# Reducing interaural tonotopic mismatch preserves binaural unmasking in cochlear implant simulations of single-sided deafness

**DOI:** 10.1101/2020.12.07.414813

**Authors:** Elad Sagi, Mahan Azadpour, Jonathan Neukam, Nicole Hope Capach, Mario A. Svirsky

## Abstract

Binaural unmasking, a key feature of normal binaural hearing, refers to the improved intelligibility of masked speech by adding masking noise that facilities perceived spatial separation of target and masker. A question particularly relevant for cochlear implant users with single-sided deafness (SSD-CI) is whether binaural unmasking can still be achieved if the additional masking is distorted. Adding the CI restores some aspects of binaural hearing to these listeners, although binaural unmasking remains limited. Notably, these listeners may experience a mismatch between the frequency information perceived through the CI and that perceived by their normal hearing ear. Employing acoustic simulations of SSD-CI with normal hearing listeners, the present study confirms a previous simulation study that binaural unmasking is severely limited when interaural frequency mismatch between the input frequency range and simulated place of stimulation exceeds 1-2 mm. The present study also shows that binaural unmasking is largely retained when the input frequency range is adjusted to match simulated place of stimulation, even at the expense of removing low-frequency information. This result bears implication for the mechanisms driving the type of binaural unmasking of the present study, as well as for mapping the frequency range of the CI speech processor in SSD-CI users.

## INTRODUCTION

The human binaural auditory system has a remarkable ability to suppress certain types of background noise. To some extent this is due to the fact that when a listener has two working ears they can pay attention to the ear with the best signal-to-noise ratio (head shadow). However, binaural intelligibility gains go beyond monaural best-ear performance when speech and noise are spatially separated (Bronkhorst and Plomp, 1988). Koenig (1950) used the term “squelch” to refer to the advantage provided by a telephone system that uses two separate microphones and delivers each signal to a different ear. This additional binaural advantage has also been referred to as binaural interaction, and binaural unmasking (Carhart et al., 1967; Bronkhorst and Plomp, 1988; Zurek, 1993; Hawley et al., 2004). Binaural unmasking has also been used more broadly to include the binaural intelligibility gains that occur under idealized listening conditions over headphones where inter-aural phase and timing relationships of target and maskers are manipulated (Carhart, 1967; Levitt and Rabiner, 1967; Bronkhorst and Plomp, 1988). One of the most surprising consequences of binaural unmasking mechanisms is that it is possible to dramatically enhance intelligibility simply by adding noise. One striking example using informational masking was provided by Freyman et al. (2001), who found that perception of a target talker in two-talker babble noise presented from the front was enhanced when the same noise was also presented from a second loudspeaker 60 degrees to the right. The additional noise signal from the right was presented 4 ms earlier than the noise signal from the front, in an attempt to engage the precedence effect. As an example, when the SNR was −4 dB and signal plus noise came only from the front, the task was very difficult and the percentage of correct responses was less than 10%. After adding the second noise source, correct responses increased to over 60%. Although not yet fully understood, the mechanisms thought to underlie binaural unmasking include peripheral, perceptual, and attentional processing of the differences and similarities in the sounds received by both ears (Bronkhorst, 2015).

One important question is: just how ‘similar’ do the left and right sounds need to be in order to elicit binaural unmasking? Most relevant for the motivation of the present study, Bernstein et al. (2015, 2016) demonstrated that it is possible to elicit binaural unmasking in listeners who receive an extremely degraded signal in the second ear. Signal plus noise was presented unprocessed to one ear, using a signal-to-noise ratio that resulted in low scores. Those scores increased significantly when a degraded version of the noise signal was also presented to the second ear. This was shown in a group of normal hearing listeners who received the second-ear noise signal after it was processed by an 8-channel noise vocoder, as well as a group of cochlear implant users with single-sided deafness (SSD-CI) who received the second-ear noise signal via direct connection to their speech processor. Both groups obtained large degrees of binaural unmasking under some conditions. For example, when the noise consisted of two talkers of the same gender as the target talker, Bernstein et al. (2016) showed that binaural unmasking for SSD-CI listeners was about 4 dB and for the normal hearing listeners in the vocoded condition it was about 7 dB. Of importance, no binaural unmasking was observed for either group when the noise signal consisted of stationary speech-shaped noise

The group of SSD-CI listeners represents a unique case to evaluate the extent to which binaural mechanisms can be restored by providing severely degraded information through a cochlear implant to an otherwise deaf (or nearly deaf) ear. Individuals with single-sided deafness (SSD) have severe-to-profound hearing loss unilaterally, e.g. PTA ≥ 70 dB HL, and normal hearing or mild hearing loss in the contralateral ear, e.g. PTA ≤ 30 dB HL (Van de Heyning et al., 2016). Despite a relatively well functioning ear, these individuals typically have deficits in sound localization and understanding speech in noise, particularly when sounds of interest are closer to the poorer ear than the better ear (Harford and Barry, 1965; Bess et al., 1986). In the past decade, cochlear implantation of the poorer ear has been increasingly adopted as a hearing intervention for SSD (e.g. Vermeire et al., 2009; Firszt et al., 2012; Zeitler et al., 2015; Van de Heyning, 2016; Arndt et al., 2017; Litovsky et al., 2019). Although originally introduced as a means of reducing tinnitus in affected SSD patients (Van de Heyning, 2009; Peter et al., 2019), it is now understood that cochlear implantation for SSD (or SSD-CI) offers many of the benefits associated with other interventions as well as some additional benefits. For example, in comparison to other interventions (such as routing of sound from the poorer to better hearing ear acoustically or through bone-conduction), cochlear implantation provides the most speech-in-noise benefit when the SNR is more favorable to the poorer ear. Additionally, cochlear implantation does not worsen performance when the SNR is more favorable to the better ear, and provides slightly improved ability to localize sound compared to other interventions (Arndt et al., 2017; Litovsky et al., 2019).

Nevertheless, levels of binaural benefit in SSD-CI users are still relatively limited in comparison to individuals with normal hearing bilaterally (Zeitler et al., 2015; Dirks et al., 2019). Adding a CI to the poorer ear only partially restores some of the mechanisms thought to underlie the advantages of binaural hearing (Arndt et al., 2011, 2017; Dirks et al., 2019; Bernstein et al., 2017). Although it is true that binaural unmasking has been demonstrated in SSD-CI listeners (Bernstein et al., 2016, 2017), it is limited to specific types of informational maskers, varies considerably across subjects, and is more modest than that experienced by normal hearing listeners. This suggests that the binaural input received by SSD-CI users is not being adequately integrated across the two ears.

Part of the problem may be due to how frequencies are mapped to electrodes in standard cochlear implant programming (Bernstein et al., 2015, 2018). At least for SSD-CI users whose unilateral hearing loss typically occurs post-lingually, a mismatch likely exists between the frequencies delivered to the CI and the cochlear locations stimulated by those frequencies prior to deafness (Landsberger et al., 2015). Although CI users can perceptually adapt to this tonotopic mismatch, their adaptation may not be complete (Sagi et al., 2010; Reiss et al., 2014; Svirsky et al., 2015; Tan et al., 2017). For SSD-CI users, tonotopic mismatch in the CI ear will create an interaural tonotopic mismatch with their normal hearing ear, and likely limit binaural processing. For example, interaural tonotopic mismatch interferes with bilateral CI users’ ability to utilize binaural cues (e.g. Kan et al., 2013, 2015).

The hypothesis that interaural tonotopic mismatch limits binaural speech-in-noise benefits in SSD-CI users has been previously examined using acoustic simulations of SSD-CI with varying degrees of interaural frequency mismatch (Zhou et al. 2017 and Wess et al. 2017). Vocoder-based acoustic simulations of cochlear implants (Blamey et al., 1984; Shannon et al., 1995) distort acoustic signals in a manner thought to approximate how sound information is processed by a cochlear implant and interpreted by an impaired auditory system. Notwithstanding limitations in accuracy of sound quality representation (Svirsky et al., 2013; Dorman et al., 2017), these models provide a platform through which to test specific hypotheses about factors that limit speech perception in various contexts.

Zhou et al. (2017) presented speech in steady-state noise to a group of normal hearing listeners under various simulated spatial configurations (i.e. over headphones). Their CI simulation incorporated different amounts of simulated electrode insertion depths while keeping the range of input frequencies constant. Under the spatial conditions tested, they found that adding the simulated CI ear generally improved speech-in-noise performance, but not for all simulated insertion depths. However, most of this binaural benefit was due to head-shadow. This result is consistent with that of Arndt et al. (2011, 2017) as well as Dirks et al. (2019). Of importance, Zhou et al. (2017) employed non-informational maskers (stationary speech-shaped noise).

Following the binaural unmasking paradigm of Bernstein et al. (2015, 2016), Wess et al. (2017) also implemented SSD-CI vocoder processing, but with varying degrees of interaural spectral and temporal mismatch. Spectral mismatch was implemented using varying degrees of simulated electrode insertion depth while keeping the input frequency range fixed (i.e., varying the mismatch between the vocoder analysis filters and the synthesis noise bands). They found that greatest binaural unmasking was achieved when vocoded maskers were aligned spectrally and temporally with unprocessed maskers presented contralaterally, and that binaural unmasking degraded substantially with increasing amounts of spectral and temporal mismatch.

The present study closely follows Bernstein et al. (2015) and Wess et al. (2017) with an important addition. The interaural frequency mismatch imposed by Wess et al. (2017) was implemented by varying simulated CI insertion depths for a fixed input frequency range. Shallower insertions produce greater interaural frequency mismatch with the contralateral normal hearing ear, especially when the input frequency range allocated to those electrodes remains fixed at clinical default settings (e.g. about 200 to 8000 Hz). In the present study, the primary premise tested is whether reducing interaural frequency mismatch by adjusting the input frequency range of the CI simulation can restore binaural unmasking in NH listeners for different simulated insertion depths. In SSD-CI users, this would be similar to adjusting the frequency map in their sound processors, which is something that can be done with existing clinical software. Providing SSD-CI users with a frequency map that minimizes tonotopic mismatch may produce better binaural unmasking benefit (a notion suggested by Bernstein et al., 2015, 2018). However, much uncertainty remains in how to accurately evaluate this mismatch in individual CI users. One advantage of conducting the present experiment with a SSD-CI simulation is that it allows us to manipulate levels of frequency mismatch with great precision and to make within-subject comparisons of different simulated electrode locations, something that is impossible with actual SSD listeners. For more shallow simulated insertion depths, minimizing interaural frequency mismatch comes at the expense of removing low frequency information. Maintaining binaural unmasking under such conditions bears importance for understanding the mechanisms driving the unmasking effect, and may help inform how to choose the best frequency-place function (i.e. frequency map) for a given SSD-CI patient.

## METHODS

### Subjects

Study participants were twenty-four adults (14 female, and 10 male) ranging in age from 22 to 44 years old (mean 28 y. o.). Prior to testing, subjects were screened for normal hearing (Grason-Stadler GSI 28). Otoscopy was used to ensure subjects’ ear canals were adequately clear of wax, and all subjects passed hearing screening, i.e. could detect pure tones of 25 dB HL at 0.5, 1, 2, and 4 kHz. Five subjects were students involved in piloting this study as part of an educational research project. IRB approval was obtained to use their data retrospectively. IRB improved informed consent was obtained prospectively for the remaining subjects.

### Stimuli

In the present study, target and masker stimuli were pre-recorded sentences drawn from the coordinated response measure (CRM) corpus (Bolia et al., 2000; Brungart, 2001a). Developed as a closed-set speech intelligibility test with masker types more relevant for military environments (Moore, 1981), CRM sentences are of the form, “Ready [call sign] go to [color] [number] now”. There are eight possible call signs (“Arrow”, “Baron”, “Charlie, “Eagle”, “Hopper”, “Laker”, “Ringo”, and “Tiger”), four possible colors (“blue”, “green”, “red”, and “white”), and eight possible numbers (1 through 8) giving 256 possible CRM sentences. The CRM corpus consists of these sentences recorded from 8 talkers (4 female and 4 male), a total of 2048 sentences. Although not a standardized test, the common structure across CRM sentences makes this test well suited to evaluate energetic versus informational masking, as well as binaural release from this masking (Brungart, 2001b; Gallun et al., 2005; Kidd et al., 2005).

In the present study, target sentences always contained the call sign “Baron” spoken by any of the male talkers in the corpus, and listeners were instructed to identify the color and number associated with the talker using this call sign. Maskers were two different, overlayed sentences spoken by any of the other male talkers in the corpus. Target and masker sentences were chosen such that when combined, each sentence used different male talkers, call signs, color, and number combinations. Same gender maskers were chosen as this combination consistently produced the greatest amount of masking and masking release in normal hearing subjects listening to unprocessed and vocoded speech (Bernstein et al., 2015), as well as in SSD-CI listeners (Bernstein et al., 2016).

For all testing conditions in the present study, unprocessed versions of target and maskers were always presented to subjects’ right ears at −3 dB SNR. That is, the two masker sentences were digitally mixed at equal root-mean-square (rms) level. The rms of the combined two-talker maskers was adjusted to be 3 dB below the rms of the target sentence, and then digitally mixed with the target. When played alone to the right ear, this monaural condition of target and maskers served as the baseline condition. For the binaural testing conditions, a vocoder-processed, or unprocessed, version of the two-talker maskers was presented simultaneously to the contralateral (left) ear. Contralateral maskers were adjusted in level to be equal in rms to the ipsilateral maskers. Prior to testing, stimuli were calibrated so that targets would be presented at 65 dBA over Etymotic ER-3A insert headphones.

### Vocoder Conditions

Binaural unmasking was evaluated with 31 binaural conditions, one where unprocessed copies of ipsilateral maskers were presented to the contralateral ear, and 30 where different eight-channel noise-band vocoder processed versions of ipsilateral maskers were presented to the contralateral ear. In vocoder processing, the acoustic input was passed through a filterbank of adjacent band-pass filters (“analysis filters”). The envelope of the output of each analysis filter was used to modulate a corresponding frequency-band of noise. The output of noise-bands were then re-synthesized to produce the vocoder processed output. As a model of how cochlear-implants process signals and delivers stimulation, the analysis filters represent the frequency range mapped to implanted electrodes and the noise-bands approximate the frequencies associated with the place of stimulation of implanted electrodes within the cochlea.

The specific vocoder processing used in the present study follows Svirsky et al. (2013). Briefly, CRM sentences were up-sampled to 48,000 samples per second and low-pass filtered to 20 kHz. A bank of sixth order bandpass Butterworth filters was then used to divide the acoustic signal into eight frequency channels. For each analysis filter, the temporal envelope was extracted by half wave rectification and third order Butterworth low pass filtering at 100 Hz. The temporal envelopes were then used to modulate a set of eight noise bands. In the present study, the 30 vocoder conditions consisted of 6 different frequency ranges for analysis filters and 5 different frequency ranges for noise-band outputs. The overall frequency ranges used for analysis filters were as follows: 63 – 6421 Hz, 188 – 7938 Hz, 313 – 9804 Hz, 438 – 12100 Hz, 563 – 14924 Hz, and 688 – 18938 Hz. The latter five of these frequency ranges were used for noise-band outputs.

Vocoder conditions are summarized in Table 1. Noise-band frequency ranges and corresponding simulated apical insertion depths are represented along the rows. Simulated apical insertion depth (in mm from the cochlear base) was calculated by applying Greenwood (1990) to the low-frequency edge of the noise bands and assuming a cochlear length of 35 mm. Frequency ranges associated with analysis filter conditions are organized along the columns. The cells for each testing condition represent the amount of basal shift, in millimeters of representative place along the cochlea (again using Greenwood, 1990), between the high-frequency edges of analysis filters and noise-bands. Possible shift conditions range from −7.5 mm (i.e. a relatively low frequency range assigned to relatively shallow electrode-array insertion) to +6 mm (i.e. a relatively high frequency range assigned to a relatively deep electrode-array insertion). The 0 mm shift conditions are the matched frequency conditions, i.e. equal analysis and synthesis frequency ranges.

**Table 1:**
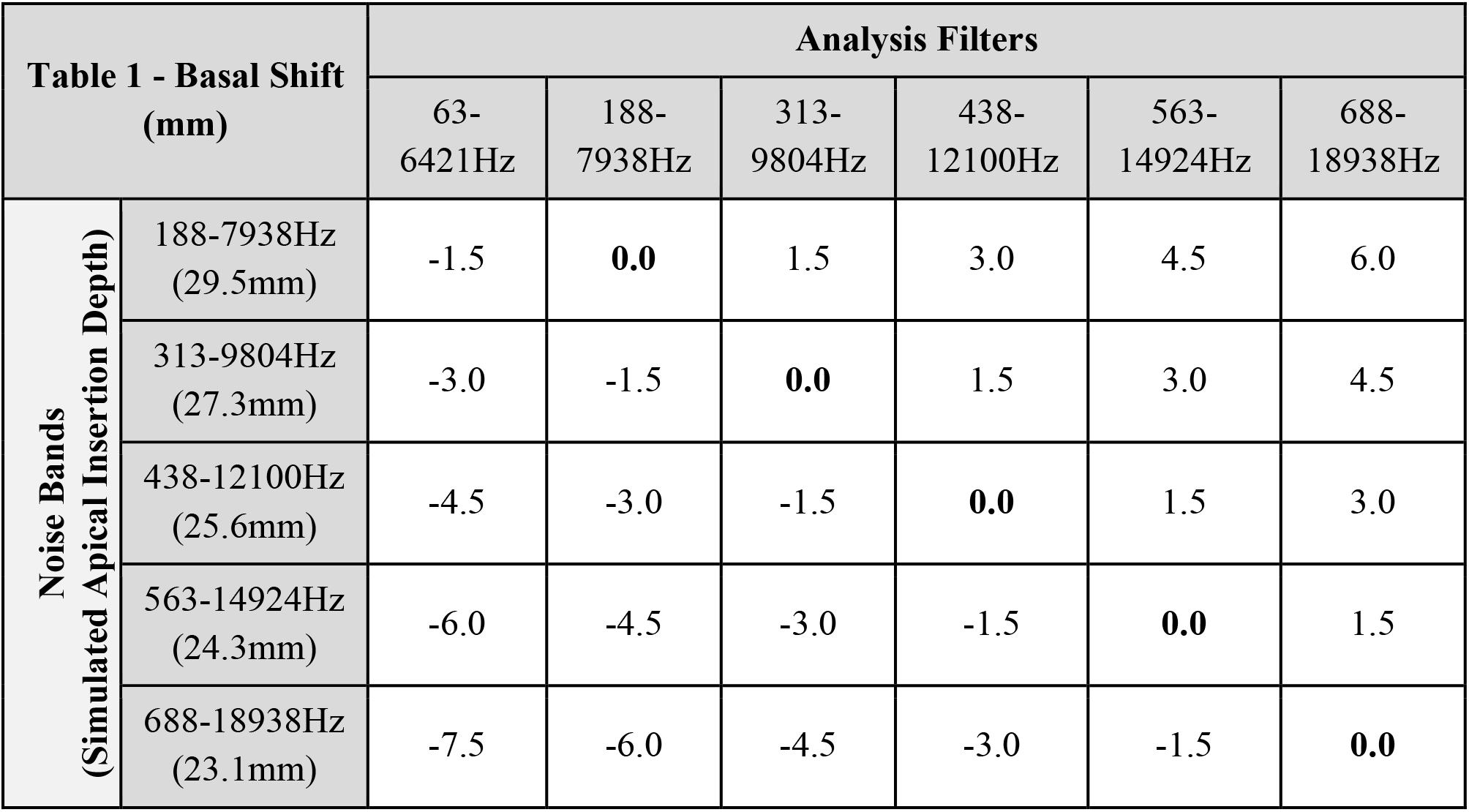
summary of frequency ranges used for analysis filters (columns) and noise bands (rows) in tested vocoder conditions. Cells represent relative frequency shift between high-frequency edges of analysis filters and noise bands, i.e. basal shift expressed in mm of cochlear place. Simulated apical insertion depth reflects cochlear place associated with lowest frequency edge of noise band conditions.

### Testing Procedure

Testing was done in two three-hour sessions. Testing conditions included the monaural condition, the binaural condition with unprocessed stimuli delivered to the contralateral ear, and the 30 binaural conditions with vocoder processed stimuli delivered to the contralateral ear. The presentation order of testing conditions was randomized for each subject. Each testing condition included 25 target and masker CRM sentences. Subjects were told to identify the color/number combination from speaker “Baron” while disregarding the two other talkers (i.e. the two-talker maskers). The number of correctly identified color and/or number combinations were evaluated as a percent correct score for each condition. Feedback regarding correct and incorrect answers was given after each trial, and subjects could view their percent correct score progressively during each condition. Ten practice trials using a binaural condition with vocoder processed stimuli were completed at the beginning of the session to confirm that subjects understood the instructions before beginning the actual task. Vocoded stimuli used for practice employed frequency ranges other than those in Table 1.

### Data Analysis

Four 2-factor Repeated Measures Analysis of Variance (RMANOVA) were carried out across different combinations of the conditions depicted in Table 1. First, a 2-way RMANOVA was conducted across all conditions of simulated insertion depths and analysis filters to assess overall differences in scores due to insertion depth, analysis filters, and any interaction between the two. Another two RMANOVAs were conducted to assess differences between conditions where analysis filters were matched to noise band filters (0 mm shift conditions) and those where analysis filters were shifted relative to noise band filters by ± 1.5 mm. Because the number of cells in Table 1 with shifts of −1.5 mm and 1.5 mm differ (5 vs 4 respectively), a separate 2-way RMANOVA was conducted for each shift condition in comparison to its corresponding matched condition for each available insertion depth. That is, the 2-way RMANOVA for the 1.5 mm shifted and matched conditions did not include the matched condition with 23 mm simulated insertion. Lastly, a 2-way RMANOVA was conducted to compare simulated insertion depth conditions with the standard clinically assigned analysis filters (2^nd^ column in Table 1) against simulated insertion depth conditions with matched analysis filters (0 mm shift). This comparison did not include the deepest simulated insertion condition (first row) because in this case the matched and clinically assigned analysis filters are equivalent.

In addition to analysis of variance two separate multiple linear regressions were conducted. One regression was used to assess the impact of using clinically assigned analysis filters on binaural unmasking for different simulated insertion depths. The other regression was similarly conducted, but when using analysis filters that matched simulated insertion depths. All five simulated insertion depth conditions were used for each regression. Analysis of Covariance was used to assess differences in regression slopes across subjects. If slopes were not significantly different, multiple regressions were performed assuming a common slope across subjects.

## RESULTS

Consistent with Bernstein et al. (2015) and Wess et al. (2017), normal hearing subjects experienced a large degree of binaural unmasking when unprocessed copies of the two-talker maskers presented to their right ears were simultaneously presented to their left ears. Average CRM percent correct scores increased from 41% in the unilateral condition (target and two-talker maskers delivered at −3 dB SNR to right ear) to 87% in the unprocessed binaural condition. A box plot of scores in the unilateral and unprocessed binaural condition is shown in Figure 1. Also consistent with those previous studies, delivering vocoder processed versions of maskers to the contralateral (i.e. left) ear produced binaural unmasking in some vocoder conditions, but to a lesser degree than the unprocessed binaural condition. Mean CRM scores (± 1 standard error) for vocoder processed binaural conditions are depicted in Table 2. Mean scores ranged from 43% to 62%.

**Figure 1:**
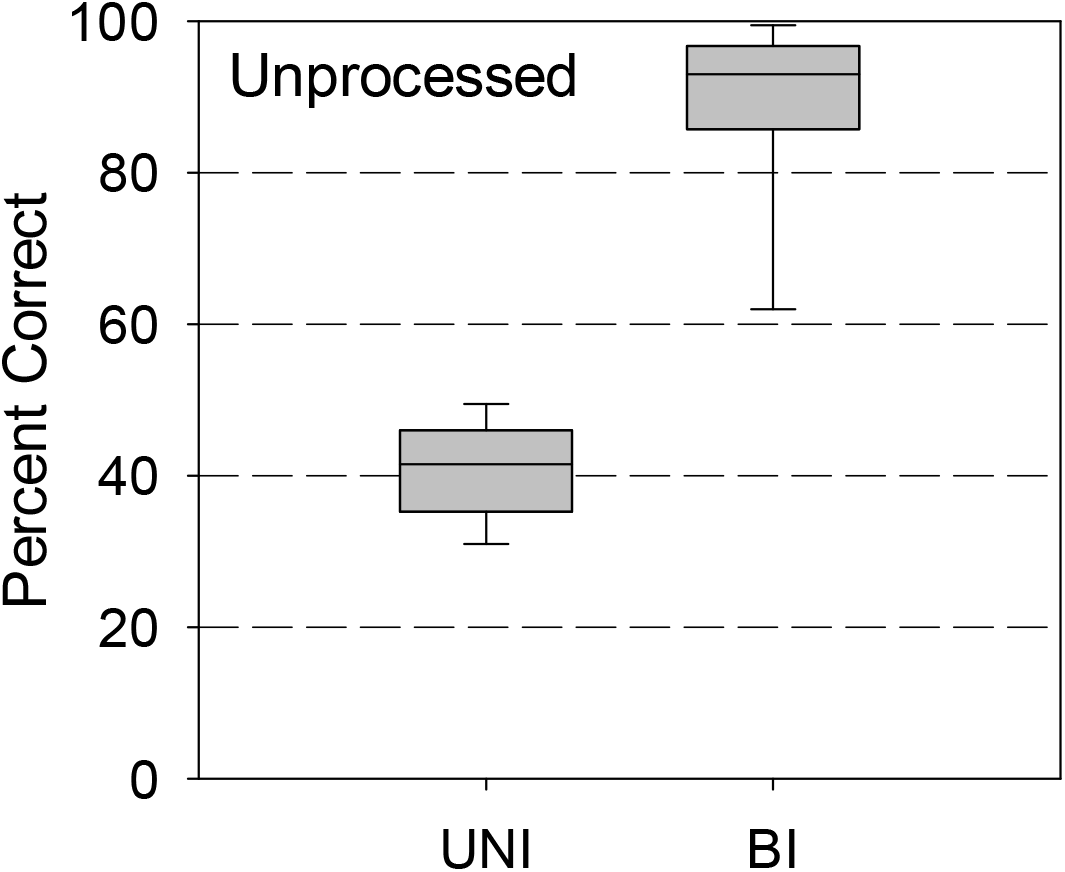
CRM percent correct scores for monaural (UNI) and binaural (BI) conditions with unprocessed stimuli

**Table 2:** CRM percent correct scores for SSD-CI binaural conditions

| Table 2 - % CRM scores (± s. e.) | | Analysis Filters | | | | | |
| --- | --- | --- | --- | --- | --- | --- | --- |
|  |  | 63-6421Hz | 188-7938Hz | 313-9804Hz | 438-12100Hz | 563-14924Hz | 688-18938Hz |
| Noise Bands<br>(Simulated Insertion Depth) | 188-7938Hz<br>(29.5mm) | 60.0<br>(2.3) | **62.4**<br>**(2.7)** | 54.8<br>(3.1) | 49.3<br>(2.1) | 48.9<br>(2.1) | 45.5<br>(2.2) |
|  | 313-9804Hz<br>(27.3mm) | 48.8<br>(2.2) | 54.2<br>(1.8) | **62.4**<br>**(3.0)** | 57.5<br>(2.0) | 53.1<br>(1.1) | 50.2<br>(2.4) |
|  | 438-12100Hz<br>(25.6mm) | 46.9<br>(2.9) | 48.7<br>(2.0) | 53.0<br>(2.2) | **59.1**<br>**(2.6)** | 54.0<br>(2.4) | 50.7<br>(2.2) |
|  | 563-14924Hz<br>(24.3mm) | 45.7<br>(2.2) | 43.0<br>(2.2) | 48.2<br>(2.2) | 55.0<br>(2.2) | **56.2**<br>**(2.2)** | 56.0<br>(2.4) |
|  | 688-18938Hz<br>(23.1mm) | 42.9<br>(1.9) | 47.6<br>(2.2) | 46.6<br>(1.8) | 49.2<br>(2.3) | 51.6<br>(2.7) | **57.5**<br>**(2.4)** |

A specific pattern emerged as to which vocoder conditions produced binaural benefit. Two-factor RMANOVA on CRM scores reveal a significant effects of analysis filters (p < 0.001) and simulated insertion depth (p < 0.001), as well as a significant interaction between the two (p < 0.001). As one can see from Table 2, mean scores are highest when the frequency range of analysis filters matched the frequency range of simulated insertion depths (diagonal entries in bold), and tend to decrease for increasing amounts of mismatch between analysis filters and simulated insertion depths. This trend is illustrated in the box plots of Figure 2 where 10%, 25%, 50%, 75% and 90% quantiles of subjects’ CRM scores are plotted as a function of relative shift (in mm) between analysis filters and simulated insertion depth for different simulated insertion depths (rows in Figure 2).

**Figure 2:**
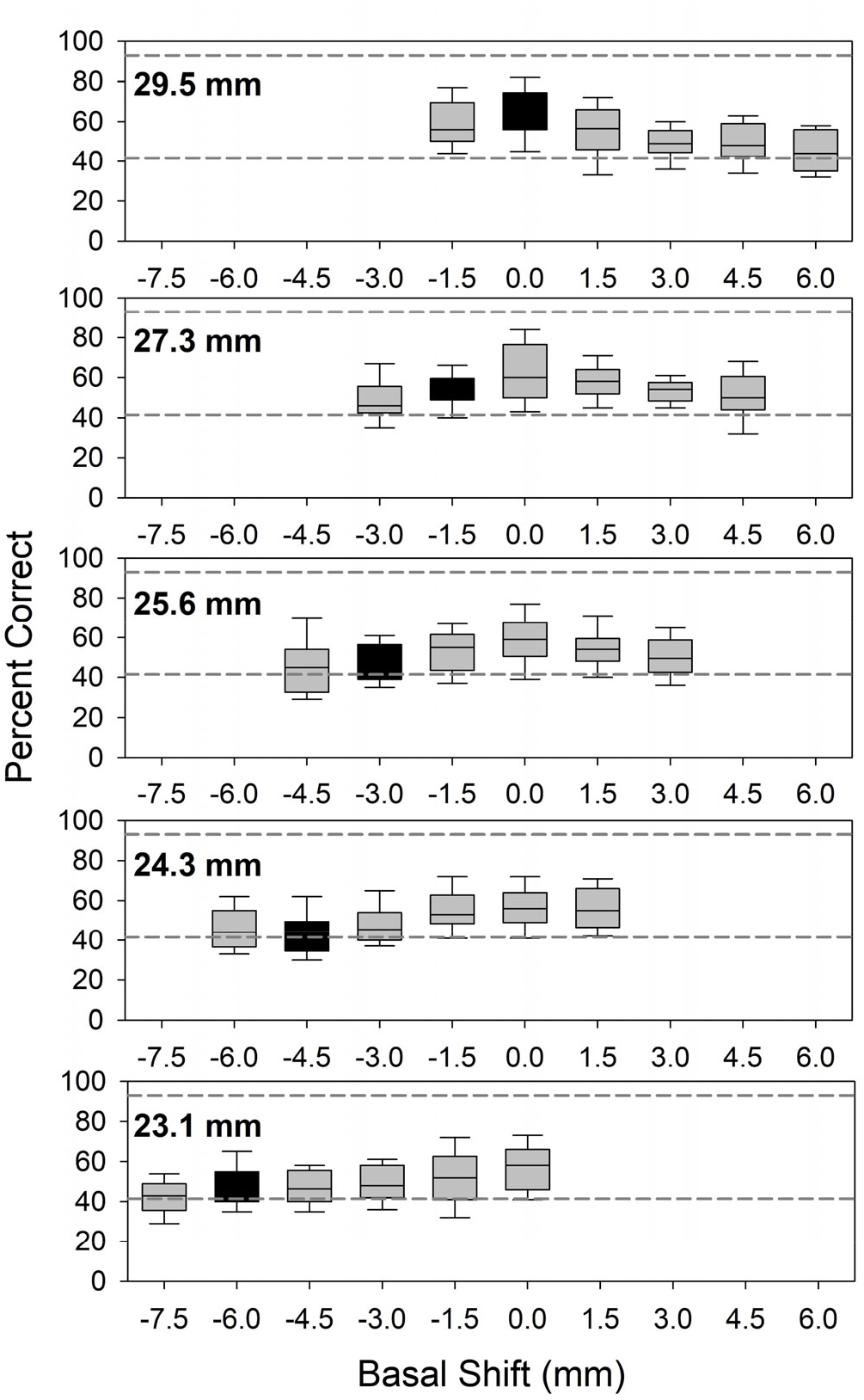
CRM percent correct scores for SSD-CI binaural conditions in terms of relative basal shift between analysis filters and simulated apical insertion depth. Black filled bars represent standard clinical frequency range.

This trend is also confirmed by separate two-factor RMANOVAs comparing shift conditions (0 mm vs −1.5 mm, and 0 mm vs 1.5 mm) across simulated insertion depths. The RMANOVA comparing 0 mm and – 1.5 mm shift conditions showed significant effects of shift (p = 0.003) and simulated insertion depth (p = 0.013), but no significant interaction (p = 0.354). Post-hoc comparisons within simulated insertion depth conditions showed that only the most extreme conditions (29 mm vs 23 mm) were significantly different (p < 0.05, holm-sidak method). The RMANOVA comparing 0 mm and + 1.5 mm shifted conditions also showed a significant effect of shift (p = 0.002), but no significant effects of simulated insertion depth (p = 0.287) or interaction (p = 0.338). Hence, the relative frequency shift between analysis filters and simulated insertion depth is the most prominent factor affecting whether vocoded contralateral maskers produce binaural unmasking.

A less prominent factor affecting binaural unmasking is the degree of simulated insertion depth, particularly for matched analysis filters. Multilinear regression on CRM scores as a function of simulated insertions depths for matched analysis filters was applied across subjects using a common slope but different intercept per subject. For matched analysis filters, CRM scores dropped by about 1% (p = 0.011) for every millimeter of shallower simulated insertion depth. Using a common slope for this multilinear regression was confirmed with an ANCOVA (i.e. slopes for individual subjects were not significantly different, p = 0.56). In contrast, simulated insertion depth did have a prominent effect on binaural unmasking when using analysis filters fixed to the typical clinically assigned frequency range (e.g. 188 – 7938 Hz for users of the Nucleus device). For standard clinical analysis filters (black bars in Figure 2), multilinear regression yields a drop in CRM scores of about 2.7% (p < 0.001) for every millimeter of shallower simulated insertion depth. ANCOVA for this multilinear regression also revealed that slopes for individual subjects were not significantly different (p = 0.15). The differential effects of clinical vs matched analysis filters on CRM scores as a function of simulated insertion depths was confirmed by a RMANOVA (p < 0.001 for matched vs standard clinical conditions; p < 0.001 for simulated insertion depth; and no interaction, p = 0.61). Hence, whereas matched analysis filters produced a 6% reduction in binaural unmasking when going from deepest to shallowest simulated insertion depths, this reduction was 16% when using a fixed analysis filter with a typical clinically assigned frequency range.

## DISCUSSION

Listeners with normal hearing can achieve substantial binaural unmasking at a target to masker ratio of −3 dB when presented with an unprocessed copy of the masker to the contralateral ear (from about 40% correct in the monaural condition to 90% correct in the unprocessed binaural condition in Figure 1). Furthermore, a significant amount of binaural unmasking remains when the contralateral ear receives a very degraded signal (from about 40% correct in the monaural condition to about 60% in the unshifted binaural conditions of Figure 2). However, this binaural enhancement is very sensitive to interaural frequency mismatch between the maskers presented bilaterally (i.e. the unprocessed maskers presented ipsilaterally with the target and the processed versions of the same maskers presented contralaterally). Even 1-2 mm of mismatch in either direction is enough to decrease binaural unmasking, and larger amounts of mismatch eliminate the effect almost completely. This result is very consistent with Wess et al. (2017), both in terms of direction and size of the effect. The present study further shows that minimizing interaural frequency shift in processed maskers largely preserves binaural unmasking, even when this is done for shallower simulated insertion depths at the expense of eliminated low frequency information (i.e. below 700 Hz). Notably, there is a reduction in the amount of binaural unmasking as low frequency information is removed, but the effect is small in comparison to the detrimental effects of interaural frequency mismatch.

The present study is intended as an analogue for understanding binaural processing in SSD-CI users, whose input to the CI is largely distorted in comparison to their contralateral ear with normal (or near-normal) hearing. That a small interaural frequency mismatch (± 1-2 mm) is enough to substantially reduce binaural unmasking in SSD-CI simulations is reminiscent of how interaural place mismatches of similar size disrupt binaural processing in bilateral CI users. Poon et al. (2009) found that JNDs for interaural time delay (ITD) in 4 bilateral CI users were poorer when interaural electrode pairs were mismatched relative to interaural electrode pairs producing minimum ITD JNDs. Their so-called ITD half-widths (i.e the interaural electrode separation producing a two-fold increase in ITD JND) spanned an average of 3 to 4 mm. Kan et al. (2013) found a similar result for ITDs in 9 bilateral CI users, although a larger interaural electrode separation tolerance was found for ILDs. Kan et al. (2013) also found that beyond ± 2 electrodes in interaural mismatch (i.e. about ± 1.5 mm in relative shift), subjects’ ability to lateralize became increasingly distorted and subjects were more likely to perceive multiple auditory images. The authors noted that distortions in lateralization due to interaural place mismatch may interfere with subjects’ ability to experience masking release, in that they may perceive physically separate sound sources as being co-located. Overall, this acute sensitivity to interaural mismatch in binaural CI hearing (whether SSD or bilateral) contrasts with the unilateral CI hearing case, where tonotopic mismatch does reduce speech perception but only for particularly large mismatches, e.g. at least 3-6 mm and greater (Svirsky et al., 2015).

Although not evaluated directly, the results of this study shed some light on an ongoing discussion regarding the mechanisms that drive binaural unmasking. Traditionally, there is a view that binaural unmasking of speech primarily operates on the lower frequency portions of the speech-signal and thus is largely attributable to ITD cues, or differences in interaural phase (Hawley et al., 2004). For example, when manipulating interaural phase of a broad-band random noise masker, Schubert and Schultz (1962) showed that binaural unmasking of speech was largely reduced in band-limited speech that removed frequencies below 800 Hz. Yet, in the present study, binaural unmasking largely persisted using simulated electrode insertion depths with matched analysis filters that removed low-frequency information below about 700 Hz. Furthermore, Wess et al. (2017) demonstrated significant binaural unmasking when vocoded maskers were spectrally matched with unprocessed contralateral maskers but presented with an interaural temporal asynchrony of −24 ms to +18 ms, i.e. temporal mismatches well beyond those at which ITD and interaural phase cues naturally operate. Taken together, these results suggest that the binaural unmasking experienced by the listeners of the present study, and those of Bernstein et al. (2015) and Wess et al. (2017), were not driven by ITD cues.

Rather, the explanation may reside in the notion of binaural unmasking by way of perceptual grouping, i.e. the segregation and streaming of sounds into distinct auditory “objects” (Bregman, 1990; Bronkhorst, 2015). Under this notion, grouping of contralaterally presented unprocessed and processed maskers into a single auditory stream provided listeners of the present study with more perceptual separation between target and maskers. One interpretation is that the improved separation of target and maskers was due to differences in perceived location. This is supported by the work of Freyman et al. (2001) wherein significant binaural unmasking was demonstrated in listeners when using the precedence effect to create the illusion of spatial separation of target and maskers presented in free field, while limiting detection advantages from true spatial separation. However, whereas the target and maskers of Freyman et al. were perceived as spatially distinct entities, those of the present study were likely perceived as a spatially distinct target versus spatially diffuse maskers due to the imperfect grouping of unprocessed and vocoded maskers, as noted by Bernstein et al. (2015).

In the present study, binaural unmasking was greatest when the frequency ranges of analysis filters and noise bands (i.e. simulated insertion depth) of vocoded maskers were matched, thus providing a vocoded copy of contralateral maskers with minimum interaural frequency mismatch. The importance of matching interaural frequency of ipsilateral and contralateral maskers to facilitate binaural unmasking is highlighted by Kidd et al. (2005). Using sine wave vocoders of the CRM corpus, target speech and speech maskers were generated with overlapping and non-overlapping frequency bands. In addition, unintelligible noise maskers with the same spectral content as speech maskers were also generated. When non-overlapping target and speech maskers were presented ipsilaterally to subjects with normal hearing, intelligibility dropped from near perfect to as low as 60% correct under the conditions of presentation level tested. However, when a noise masker spectrally matched to the speech masker was added contralaterally, subjects’ intelligibility recovered to just over 80% correct. That is, substantial binaural unmasking was achieved by providing unintelligible noise to the contralateral ear, albeit spectrally matched to the ipsilateral speech masker. This result of Kidd et al. demonstrates the importance of matched interaural frequency as an effective cue or heuristic to affect perceptual grouping and facilitate the perceptual separation of target and masker.

An important caveat to the present study is that its results were achieved using informational masking, and the release thereof (Brungart et al., 2001; Freyman et al, 2001; Gallun et al., 2005; Kidd et al., 2005). Indeed, Bernstein et al. (2015) has demonstrated little to no binaural unmasking when masker types were stationary or modulated noise, or even when maskers were interfering talkers of different gender than the target. This result is also consistent with the other studies cited in the introduction of the present study that showed lack of squelch in SSD-CI acoustic simulations and in SSD-CI users when using energetic masking (Zhou et al., 2017; Bernstein et al., 2016, 2017; Dirks et al., 2019). The specificity of this effect to informational masking is one indication that this type of unmasking relates more to perceptual grouping, something thought to occur at a relatively higher stage of auditory processing that interacts with attention (Bronkhorst, 2015). Indeed, binaural release from informational masking is relatively non-specific to ITD or ILD cues, occurring over a wide range of values for either cue, such that similar amounts of unmasking occur whenever the perceived location of target and masker differ (Gallun et al., 2005). Under this account of binaural unmasking, the present study demonstrates the sensitivity of perceptual grouping to interaural frequency mismatch, and the ability to restore binaural grouping of maskers by minimizing this mismatch, albeit at the expense of removing low frequency information.

Given the relative intolerance of binaural hearing with CIs to interaural tonotopic mismatch, there may be clinical benefit to reducing or minimizing this mismatch (Bernstein et al., 2015; Wess et al., 2017). This is particularly true for SSD-CI users who could potentially benefit more from binaural advantages, if any, offered by their CI ear. The present study shows that given an amount of binaural frequency mismatch, reducing it by modifying analysis filters is sufficient to recover much of the lost binaural enhancement effect. The implication is that modifying a CI speech processor’s frequency allocation map may be an option to reduce interaural tonotopic mismatch in SSD-CI subjects and possibly restore some degree of binaural enhancement beyond those due to head shadow (or better ear listening). Although such a modification comes at the expense of removing low-frequency information, especially for shallower electrode insertions (Faulkner et al., 2006), the present study suggests removing low frequency information may only have a modest effect on binaural unmasking in SSD-CI listeners if one can thereby minimize interaural tonotopic mismatch.

## Notes

### Competing Interest Statement

The authors have declared no competing interest.

### Summary of Updates

Article has been submitted to The Journal of the Acoustical Society of America. After it is published, it will be found at http://asa.scitation.org/journal/jas.

